# CRISPR/Cas9-mediated generation of a *de novo C9ORF72* knock-in isogenic iPSC cell bank to model ALS and FTD

**DOI:** 10.1101/2025.11.12.687564

**Authors:** Niamh L. O’Brien, Erin Hedges, Remya R. Nair, Alexander J. Cammack, Mireia Carcolé, Deniz Vaizoglu, Sara Tacconelli, Katie Sutherland, Chloe L. Fisher-Ward, Ali Raza Awan, Simon Topp, Alfredo Iacoangeli, Adrian M. Isaacs, Thomas J. Cunningham, Marc-David Ruepp, Sarah Mizielinska

## Abstract

**Background:** The pathogenic G_4_C_2_ repeat expansion in the *C9ORF72* gene is the most common genetic cause of amyotrophic lateral sclerosis (ALS) and frontotemporal dementia (FTD). Studies focused on delineating the underlying perturbed mechanisms resulting from this genetic mutation are often confounded by the heterogeneity present in current disease models, such as patient-derived iPSC lines, with estimations of up to 50% of the variation in iPSC cell phenotypes resulting from inter-individual differences. Isogenic models, in which the pathogenic mutation is introduced into a defined genetic background, offer a powerful approach to isolating mutation-specific effects and enable high-resolution comparison across distinct ALS/FTD-associated mutations. Such models are essential for uncovering convergent disease mechanisms and improving reproducibility in ALS/FTD research.

**Methods:** A two-step scarless CRISPR/Cas9 genome editing strategy was used to generate isogenic human iPSC lines carrying a *de novo* knock-in of a disease-length G_4_C_2_ repeat expansion in the *C9ORF72* locus. The resulting lines underwent thorough quality control and were differentiated into lower motor neurons and assessed for the presence of key ALS/FTD pathologies, including changes to *C9ORF72* mRNA and protein expression, RNA foci and dipeptide repeat proteins.

**Results:** Two *C9ORF72* knock-in iPSC lines were generated with 631 and 600 G_4_C_2_ repeats, alongside an isogenic genome editing control line. The *C9ORF72* G_4_C_2_ repeat expansion knock-in iPSC lines exhibit both loss-of-function and gain-of-function pathological features characteristic of ALS/FTD. Compared to the parental wild-type KOLF2.1J line and isogenic (wild-type) CRISPR control line, these exhibit a significant reduction in *C9ORF72* mRNA and protein levels, the presence of RNA foci accumulation, and a marked increase in poly(GA) and poly(GP) dipeptide repeat protein levels in iPSCs and motor neurons.

**Conclusions:** This is one of the first reports of a successful knock-in of the pathogenic *C9ORF72* G_4_C_2_ repeat expansion into a human iPSC line, establishing a genetically defined and physiologically relevant model of ALS/FTD. These isogenic lines recapitulate both key loss- and gain-of-function disease pathologies, providing a crucial complement to existing patient-derived iPSC banks. By eliminating confounding genetic background variability, these cell lines will enable more precise interrogation of *C9ORF72*-linked pathomechanisms and offer a robust platform for comparative studies across the ALS and FTD spectrum, mechanistic investigations, and future therapeutic targeting with enhanced translational relevance.

## Introduction

Amyotrophic lateral sclerosis (ALS) and frontotemporal dementia (FTD) are fatal neurodegenerative diseases characterised by selective loss of distinct types of neurons: predominantly motor neurons in ALS and cortical neurons in FTD. An intronic GGGGCC (G_4_C_2_) hexanucleotide repeat expansion (HRE) in the *C9ORF72* gene is the most common genetic form of ALS and FTD, accounting for up to 35% of familial ALS^1,2^ and 25% of FTD cases^3^.

There are currently three non-exclusive proposed mechanisms through which the presence of the HRE leads to the degeneration of neurons. Firstly, a loss-of-function mechanism has been proposed whereby the presence of the repeat expansion leads to reduction of the endogenous *C9ORF72* mRNA and protein^4–6^. The presence of the repeat expansion can also exert toxic gain of function mechanisms, as the presence of the repeat results in the transcription of RNA containing both sense and antisense repeat sequences^1,7,8^ and the production of repeat encoded dipeptide repeat proteins (DPRs)^9–11^. Both sense and antisense RNA foci are detected in postmortem tissue from individuals with ALS and FTD^1,12^. Unconventional repeat associated non-ATG dependent (RAN) translation of the *C9ORF72* HRE RNA sense strand results in the production of the DPR polypeptides glycine-alanine (GA), glycine-arginine (GR), and glycine-proline (GP), and translation of the antisense strand produces the polypeptides proline-arginine (PR), proline-alanine (PA), and proline-glycine (GP)^13^. These DPRs have been found to aggregate in patient brain and spinal cord neurons^7,14^ with the arginine-containing poly(GR) and poly(PR), and poly(GA) known to have the most neurotoxic potential^15–19^. A wealth of cellular mechanisms have been demonstrated to link the aforementioned loss- and gain-of-function entities to neurodegeneration^20^, thus using robust models that display the full range of molecular pathologies is crucial for studying *C9ORF72* related diseases.

Induced pluripotent stem cells (iPSCs) are important tools for both discovery biology and therapy development. They offer the opportunity to model specific disease relevant phenotypes in human cells affected by neurodegenerative conditions such as ALS and FTD and are proven to be robust models for furthering our understanding of disease progression. Isogenic stem cell banks offer the ability to control for both genetic and epigenetic heterogeneity therefore reducing the variability in results. Genomic profiling of a large cohort of iPSC lines by the Human Induced Pluripotent Stem Cells Initiative (HipSci) have reported that 5-46% of phenotypic variability can be attributed to genetic differences between individuals^21^. Compared to patient-derived iPSC lines, which, by nature, have diverse genetic backgrounds, genome-edited isogenic models provide a controlled platform to study disease phenotypes attributable solely to specific mutations, in this case the *C9ORF72* mutation. This approach enhances reproducibility, enables precise mechanistic studies, and will facilitate direct comparison with other ALS/FTD-linked mutations, such as ALS mutations in *FUS* and *TARDBP* engineered on the same iPSC genetic background^22^.

The goal of this study was to insert a large repeat expansion into the *C9ORF72* endogenous locus on the well-established KOLF2.1J background^22^ as a key new tool for ALS and FTD research. Here we demonstrate the successful generation of two isogenic *C9ORF72* iPSC lines with disease-length HRE which display all major loss- and gain-of-function *C9ORF72* ALS/FTD related pathological features. The resultant iPSC bank offers a powerful tool for downstream applications including therapeutic targeting, transcriptomic profiling, and investigation of early-stage disease processes for the most common genetic cause of ALS and FTD.

## Results

### Generation of a C9ORF72 knock-in iPSC line

The approach for inserting the HRE into the endogenous *C9ORF72* genomic locus of the KOLF2.1J iPSC line is outlined in Figure 1A. A two-step process was used by first generating an intermediate landing pad iPSC line, which enabled the use of a highly efficient and specific gRNA target site, in heterozygosity, to minimise undesired gene editing. This was followed by insertion of the HRE using the *de novo* gRNA target site. The *C9ORF72* HRE donor plasmid used is a previously described pJazz linear plasmid^23^ engineered to contain 1027 G_4_C_2_ repeats flanked by 2 kilobases of homology upstream and 3 kilobases downstream of the repeat locus, respectively. Additionally, the plasmid contains a positive selection cassette, expressing mNeonGreen and conferring puromycin resistance, flanked by PiggyBac transposase cut sites (Supplementary Figure 1A) to allow for the seamless excision of the cassette following selection of iPSC colonies positive for the integration of the HRE.

**Figure 1.**
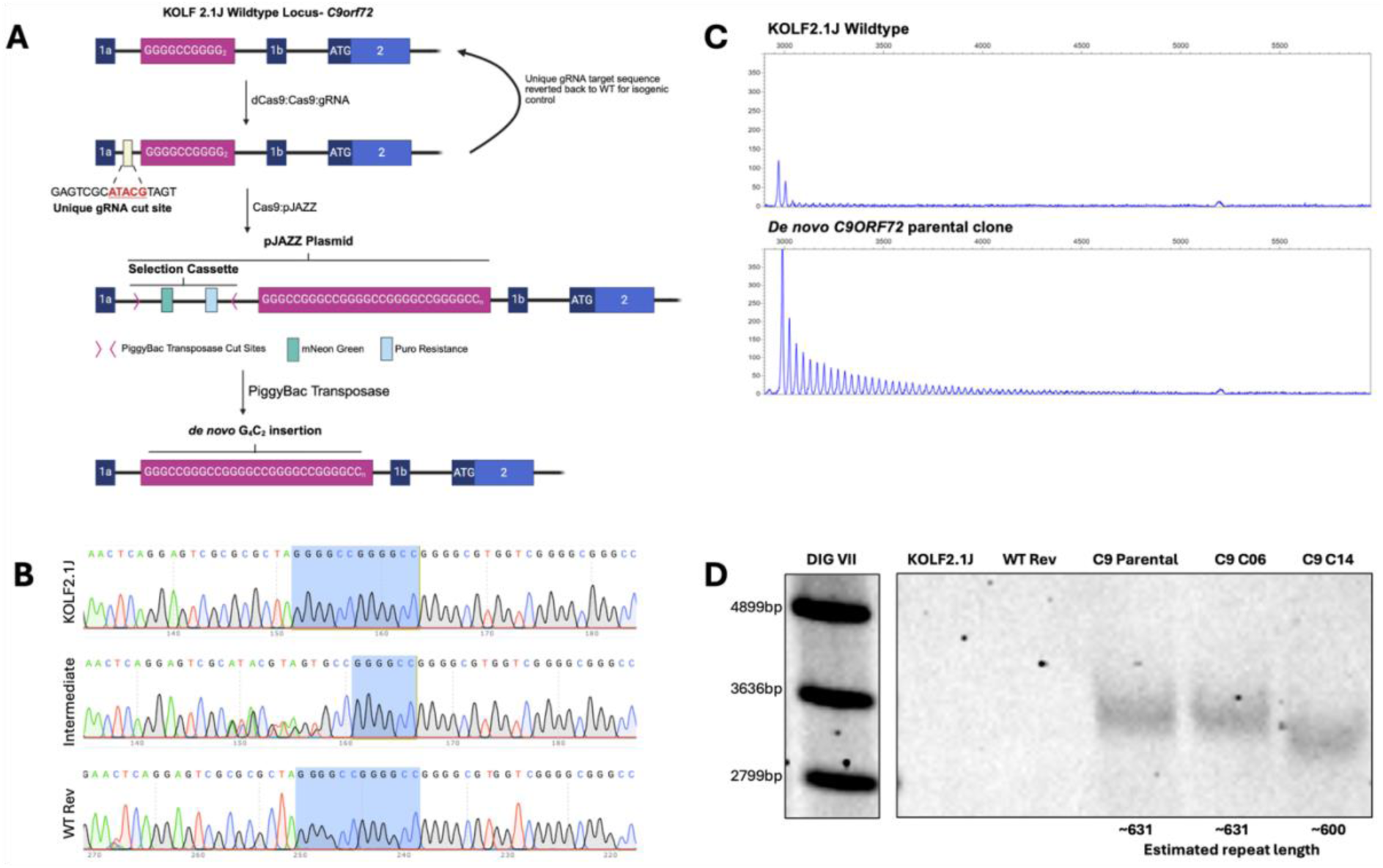
CRISPR/Cas9 genome editing insertion of the HRE at the endogenous *C9ORF72* locus. (A) Overview of the CRISPR/Cas9 strategy for insertion of the HRE. (B) Validation of the intermediate line showing the introduction of the *de novo* heterozygous gRNA target site and the subsequent reversion of this insertion back to the WT sequence for the wild-type CRISPR control (WT Rev). (C) Expanded G_4_C_2_ repeats were detected in the parental clonal lines prior to subcloning with repeat prime PCR. (D) Southern blot demonstrating the presences of a repeat expansion in the *C9ORF72* parental (∼631 repeats) and the subcloned lines, C06 (∼631 repeats) and C14 (∼600 repeats), following the removal of the selection cassette compared to the KOLF2.1J and WT Rev CRISPR control.

In the first step, point mutations were identified that could be inserted upstream of the two G_4_C_2_ repeats present in the KOLF2.1J line at the *C9ORF72* locus to produce a highly efficient novel gRNA target site otherwise not present in the human genome. Designing a target site for the gRNA with high on-target and low off-target cleaving scores was also prioritised. This intermediate iPSC line was used to generate both the *C9ORF72* mutant HRE and the wild-type revertant CRISPR control (WT Rev) lines (Figure 1A & B). The WT Rev line was generated as the CRISPR control, undergoing the same number of CRISPR experiments with the same gRNA as the resulting *C9ORF72* HRE knock-in lines. PCR and Sanger sequencing confirmed the reversion of the introduced gRNA target site back to the endogenous *C9ORF72* sequence (Figure 1B). All single-stranded oligodeoxyribonucleotide (ssODN) template oligonucleotides and gRNA sequences are listed in Supplementary Table 1.

In parallel, the intermediate line was used to insert the pJazz plasmid containing 1027 G_4_C_2_ repeats into the *C9ORF72* locus. Following CRISPR/Cas9, iPSC colonies were selected for screening based on their resistance to puromycin and expression of nuclear mNeonGreen signal. Repeat prime PCR (rpPCR) confirmed the presence of a repeat expansion in several iPSC clones (Figure 1C). Correct insertion of the repeat expansion was validated using primers targeting outside of the homology region and into the selection cassette (Supplementary Table 2). The positive selection cassette was removed from the parental lines using a PiggyBac transposase excision-only plasmid^24,25^.

This resulted in two iPSC lines with the repeat expansion present and the cassette removed: C06 and C14. Southern blotting was used to determine the successful insertion of approximately 631 G_4_C_2_ repeats in the parental *C9ORF72* line, prior to removal of the positive selection cassette. Following cassette excision, repeat expansions were detected in two subclones, with clones C06 and C14 harbouring 631 and 600 repeats respectively, which were absent in the KOLF2.1J wild-type and WT Rev control (Figure 1D, Supplementary Figure 2A). To further validate the repeat sizing and confirm correct on-target integration, Nanopore adaptive sequencing targeting the *C9ORF72* gene region was undertaken, specifically targeting chr9:27543537-27603536. Nanopore sequencing sized the insertion in C06 as 700 repeats, with C14 containing up to 500 repeats (Supplementary Figure 1B), broadly matching the results from the Southern blot (Figure 1D, Supplementary Figure 2A). Illumina paired-end whole genome sequencing also confirmed the presence of the HRE in heterozygosity in the two intended lines and excluded the presence of any off-target plasmid integration sites in all three lines. SNP array karyotyping confirmed a normal karyotype of 46;XY for the *C9ORF72* HRE knock-in, WT Rev control and KOLF2.1J iPSC lines (Supplementary Figure 3A) and the absence of off-target indels in each of the CRISPR generated lines (Supplementary Figure 3B).

### *C9ORF72* HRE knock-in lines express iPSC markers and can efficiently differentiate to lower induced motor neurons

Next, the quality of the engineered iPSC lines was assessed to ensure a lack of inherent differences in pluripotency or differentiation capabilities. At the iPSC level, all cell lines expressed similar levels of the pluripotency markers TRA-1-81, Nanog and SSEA-4 by immunostaining (Figure 2B) and *OCT3/4* by qPCR (Figure 2E). To ensure that cells maintained the ability to differentiate into motor neurons, iPSCs were differentiated into lower induced motor neurons (liMNs) using small molecule patterning combined with doxycycline induced-*NGN2* expression (Figure 2A)^26^. At 28 days *in vitro C9ORF72* HRE knock-in and control lines robustly expressed the neuronal marker MAP2 (Figure 2C) and the motor neuron specific marker ISLET1 at similar levels (Figure 2C&D). Additionally, qPCR showed similar levels of *MNX1* (*HB9*) expression across all cell lines, a key marker of spinal motor neuron identity, which was significantly increased compared to expression in iPSCs (Figure 2F).

**Figure 2:**
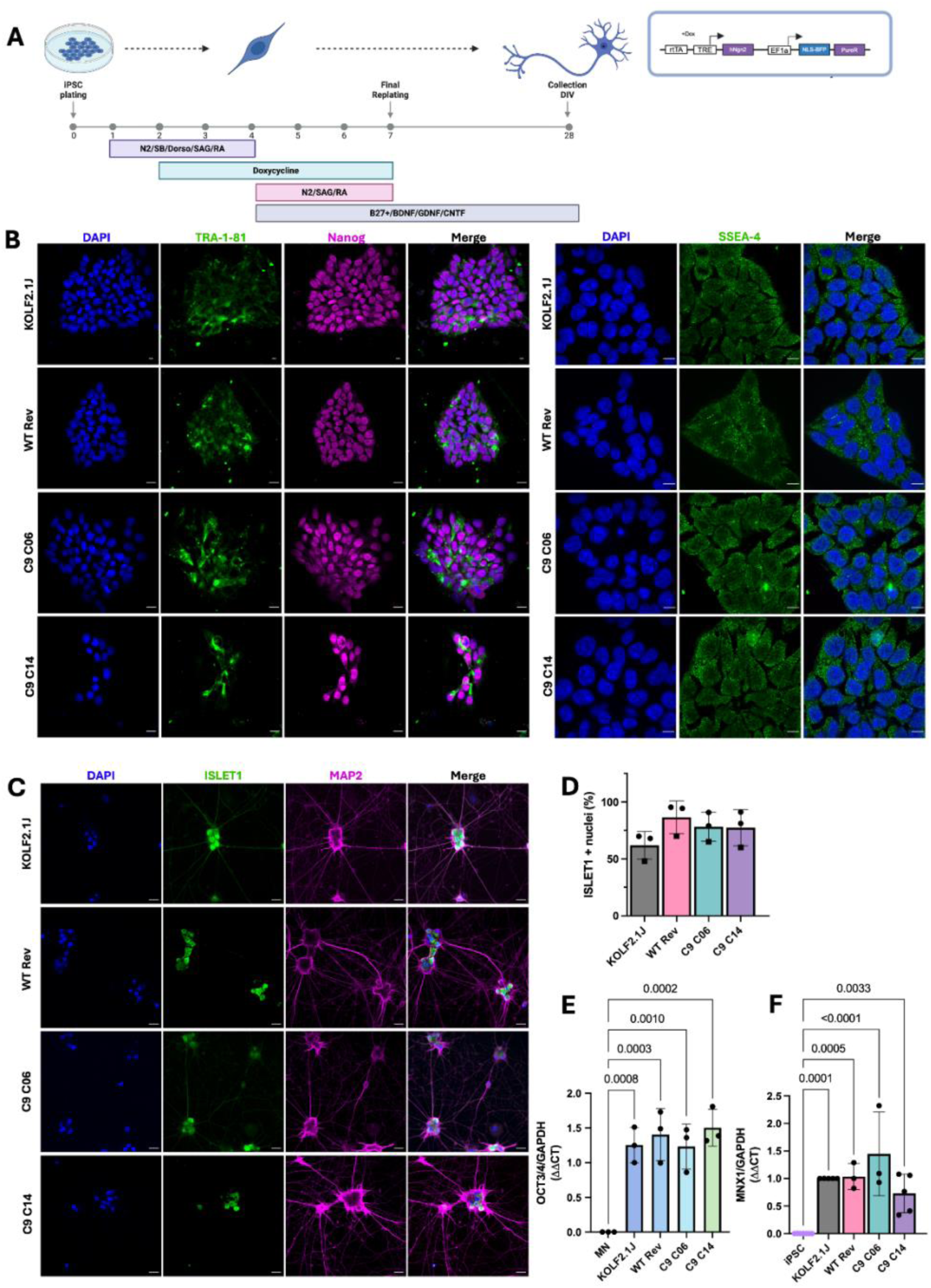
The *C9ORF72* HRE knock-in and control iPSC lines express iPSC markers and differentiate efficiently into lower induced motor neurons. (A) Protocol overview for the differentiation pipeline for generation of liMNs. (B) Maximum intensity projections of iPSCs immunolabelled for the iPSC markers TRA-1-81, Nanog (scale bars 25mm) and SSEA-4 (scale bar 10mm). (C) Gene edited iPSCs differentiate to MAP2 and ISLET1 positive liMNs with the same efficiency as the origin KOLF2.1J line (scale bars 25mm). (D) All lines have similar expression of the motor neuron marker ISLET1 at DIV 28. *n*=3 biological replicates, 2 technical replicates per differentiation. (E) qPCR data shows an increase in *OCT3/4* expression in the iPSC lines compared to DIV 28 liMNs. (F) A significant increase in the motor neuron marker *MNX1* was observed at DIV 28 compared to iPSCs. Bars are mean ± standard error, *n*=3-5 biological replicates. Samples were analysed for all experiments using a one-way ANOVA with post hoc Šídák’s multiple comparisons test with comparison versus the wild-type controls.

### *C9ORF72* HRE knock-in lines display a haploinsufficiency phenotype

One of the pathological features most commonly reported in *C9ORF72*-ALS/FTD patient post-mortem tissue and patient derived iPSC lines is the reduction of up to 50% of both total *C9ORF72* mRNA and protein levels^1,27,28^. Different transcriptional start sites in the *C9ORF72* gene lead to the generation of three main mRNA variants encoding two isoforms of the C9ORF72 protein, where the HRE is located either in the promoter region or the first intron followed by use of non-coding exon 1b (Figure 3A). Interestingly, although total levels of *C9ORF72* mRNA are found to be reduced, levels of intron 1a, upstream of the HRE, are increased in *C9ORF72*-ALS/FTD^29^. Using previously described primers^30–32^, expression of different *C9ORF72* variants was quantified, targeting total *C9ORF72* expression (all variants), variant 2, and intron 1a-containing mRNA (Figure 3A). Total *C9ORF72* expression levels were significantly decreased in C14 iPSCs (*p*<0.03), with a trend towards reduction in C06. No significant difference in variant 2 or intron 1a expression was detected in our knock-in iPSC lines compared to controls (Figure 3B). Similar trends were observed in liMNs with a significant reduction in total *C9ORF72* in C14 (*p*=0.01) and 20-25% reduction trend in C06, no significant change in variant 2, and a trend towards increase in intron 1a expression in *C9ORF72* HRE knock-in lines compared to controls (Figure 3C).

**Figure 3.**
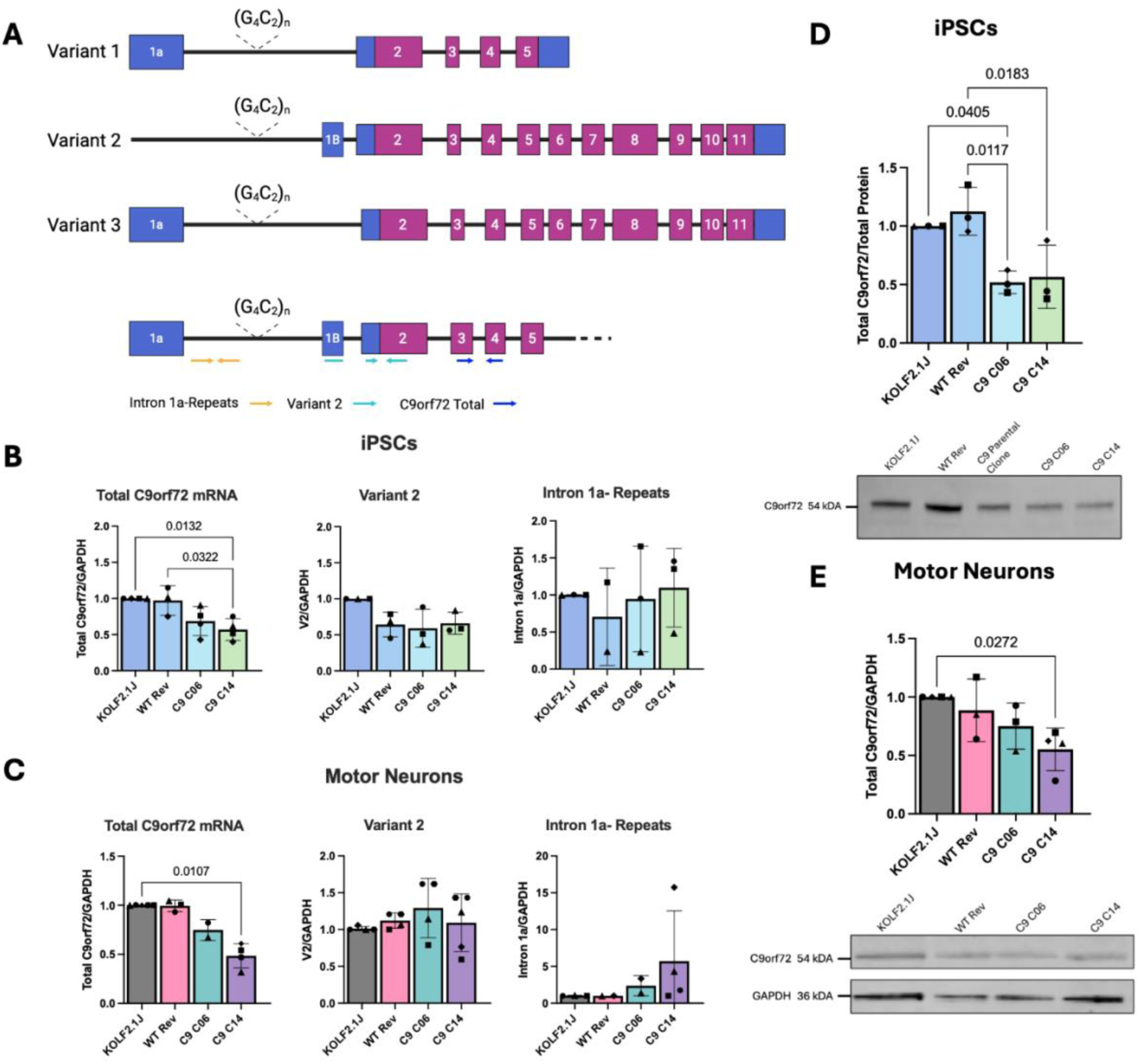
*C9ORF72* HRE knock-in lines display haploinsufficiency. (A) Three main variants of *C9ORF72* and the target sites of used primers. (B) qPCR of *C9ORF72* variant levels in iPSCs demonstrates a significant reduction of total *C9ORF72* mRNA in C14 with a trend towards a reduction in C06. No significant difference was observed in the levels of variant 2 (the major isoform) or intron1a containing transcripts. (C) qPCR from DIV 28 liMNs shows a significant reduction in total *C9ORF72* in C14. No significant difference in the expression levels of variant 2 or intron 1a were observed at this time point. (D&E) Western blot analysis of total C9ORF72 protein shows a 25-50% reduction in both C9ORF72 knock-in (D) iPSC and (E) liMNs compared to the wild-type controls. n=2-4 independent differentiations per line. Samples were analysed for all experiments using a one-way ANOVA with post hoc Šídák’s multiple comparisons test with comparison versus the wild-type controls

Western blot analysis of total C9ORF72 protein demonstrates a 50% reduction in both C06 (*p*<0.05) and C14 iPSCs (*p*<0.02), with a smaller reduction in DIV 28 liMNs of 25% and 50%, respectively, compared to the wild-type controls (Figure 3D&E, Supplementary Figure 2B&C).

### Knock-in *C9ORF72* HRE iPSC derived motor neurons exhibit both sense and antisense RNA foci

The presence of both sense and antisense HRE RNA foci is a key gain-of-function pathological feature of *C9ORF72* related ALS and FTD^33^. Typically, RNA foci are found abundantly in neurons, and have also been detected in glial cells such as astrocytes and microglia^8,34^.

Knock-in cells were differentiated to liMNs and levels of RNA foci measured using fluorescent *in-situ* hybridization at 28 days *in vitro* (DIV). The two *C9ORF72* HRE lines both clearly demonstrate the presence of both sense and antisense HRE RNA foci, which are absent in wild-type KOLF2.1J and WT Rev control lines (Figure 4A&B). Foci were detected in the nucleus and cytoplasm of neurons, with foci-positive neurons often containing multiple foci per cell. Quantification of RNA foci load demonstrated an increase in the number of neurons containing sense foci in both *C9ORF72* clonal lines and a significantly higher number of neurons containing antisense foci compared to control lines (p<0.05) (Figure 4B), recapitulating RNA foci burden present in postmortem brain tissue from individuals with *C9ORF72* ALS/FTD^8,35,36^.

**Figure 4.**
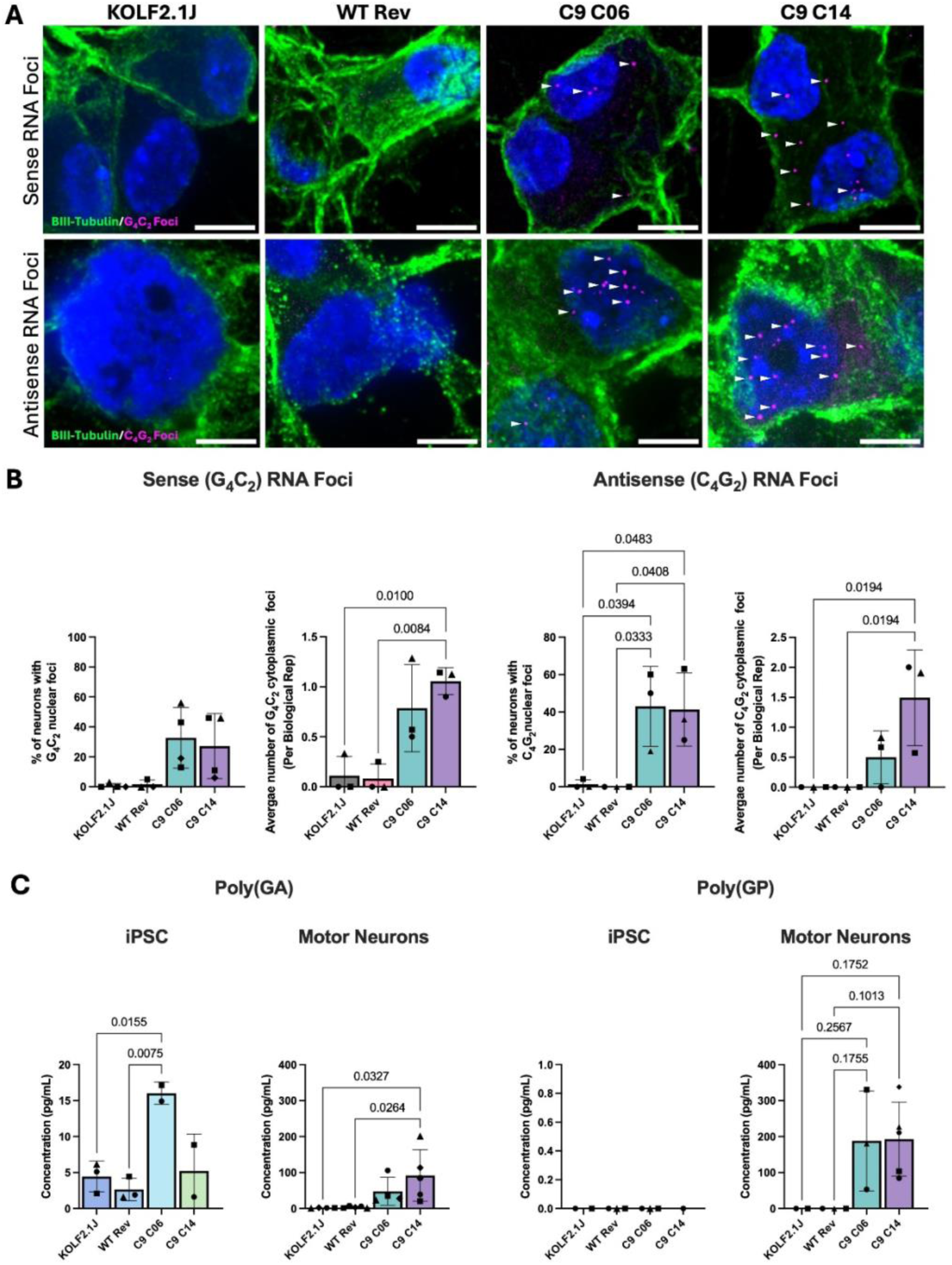
*C9ORF72* iPSC derived liMNs (DIV 28) contain both sense and antisense RNA foci and express DPRs. (A) Maximum projection of DIV 28 liMNs show increased abundance of sense and antisense nuclear and cytoplasmic HRE foci in knock-in but not control neurons (n=3, 2 technical replicates per differentiation, 5-10 images acquired per well, scale bar 10μm). (B) Knock-in lines had a significantly higher burden of sense foci and a significant increase in antisense foci compared to the wild-type control liMNs. There was also an enrichment of sense and antisense cytoplasmic foci (n=3, 2 technical replicates per differentiation, 5-10 images acquired per well, scale bar 10µm). (C) MSD analysis of the *C9ORF72* HRE lines shows a significant increase in poly(GA) at both the iPSC and liMN stage. Poly(GP) was not detected in the iPSC lines but was enriched in the *C9ORF72* liMNs at DIV 28. Samples were analysed for all experiments using a one-way ANOVA with post hoc Šídák’s multiple comparisons test with comparison versus the wild-type controls

### DPR detection in *C9ORF72* HRE knock-in iPSC lines

The levels of DPRs in the *C9ORF72* knock-in lines were measured by immunoassay for the sense DPR poly(GA) and the sense or antisense DPR poly(GP). Poly(GA) is detected in *C9ORF72* HRE lines at both the iPSC and liMNs stage, though concentration is dramatically higher in liMNs (Figure 4C). This reached significance for C06 in iPSCs and C14 in liMNs (*p<*0.05) compared to the KOLF2.1J and WT Rev control. For poly(GP), there were no detectable levels in *C9ORF72* HRE iPSCs, but liMNs demonstrate a clear increase in both *C9ORF72* lines compared to the control lines (Figure 4C). Together, loss of C9ORF72 protein and detection of RNA foci and DPRs confirms that our novel *C9ORF72* HRE knock-in iPSC lines demonstrate both loss- and gain-of-function molecular phenotypes.

## Discussion

A pathogenic G_4_C_2_ repeat expansion in the *C9ORF72* gene is the most common genetic cause of ALS and FTD, yet studies focused on delineating the underlying perturbed mechanisms are often confounded by the heterogeneity present in current disease models, such as patient-derived iPSC lines. The aim of this study was to develop a *de -novo* isogenic iPSC line, where the HRE was inserted into the *C9ORF72* gene on the background of the well-established wild-type KOLF2.1J line^22^. A high efficiency two-step CRISPR/Cas9 approach was designed that allowed the seamless integration of over 600 G_4_C_2_ repeats into the endogenous *C9ORF72* locus, which generated two *C9ORF72* HRE knock-in iPSC lines. Additionally, an isogenic CRISPR wild-type revertant line was generated which underwent the same number of genomic editing experiments as the isogenic *C9ORF72* HRE lines, allowing to control for experimental variation while keeping cell line passage numbers similar, which is essential for robust comparability. These lines have undergone stringent quality control checks to ensure the absence of genomic abnormalities, including karyotype alteration or introduction of on- or off-target mutations, and the maintenance of pluripotency and differentiation capacity.

The generated knock-in *C9ORF72* HRE lines display key ALS/FTD phenotypes mirroring pathology observed in both patient iPSCs and derived neurons, and post-mortem tissue. These lines exhibit a reduction in *C9ORF72* mRNA and protein levels at the iPSC and motor neuron stage, demonstrating a significant haploinsufficiency phenotype. They also accumulate both nuclear and cytoplasmic sense and antisense HRE RNA foci in motor neurons derived from the *C9ORF72* mutant lines compared to their wild-type controls, as well as the DPR products poly(GA) and poly(GP), recapitulating both RNA and protein toxic gain-of-function pathologies associated with *C9ORF72* ALS and FTD. Targeting HRE RNA in *C9ORF72* ALS/FTD has been a major therapeutic strategy given their central role in disease pathogenesis. However, to date, trials focusing on antisense oligonucleotide (ASO) interventions, such as the BIIB078 ASO trial ^37^ and the WVE-004^38^, have not passed their primary endpoint, either failing to show clinical benefit or showing functional decline at higher doses in the absence of a clinical benefit. Specifically, the BIIB078 trial targeted the ASO to the sense G_4_C_2_ strand to reduce sense strand derived DPRs and RNA foci, suggesting that for beneficial therapeutic development antisense repeat pathologies may also need to be targeted^39^.

The development of this isogenic *C9ORF72* iPSC bank is critical for enabling reproducible high-resolution determination of mutation-specific alterations in the most common form of ALS and FTD, and to identify convergent and divergent disease mechanisms and phenotypes through comparison across the ALS-FTD spectrum. Indeed, this cell bank will integrate with an independent effort to generate an isogenic *C9ORF72* iPSC line led by Coneys and colleagues^40^, and also the publicly available iPSC bank generated by the Induced Pluripotent Stem Cell Neurodegenerative Disease Initiative (iNDI), which all use the KOLF2.1J background. Furthermore, comparison to patient iPSC lines may reveal additional genetic drivers of disease heterogeneity within *C9ORF72* cases.

In summary, by eliminating confounding genetic background variability, this *C9ORF72* isogenic iPSC bank enables precise interrogation of *C9ORF72*-linked pathomechanisms and comparative studies to address inter- and intra-disease heterogeneity, offers a robust platform for therapeutic assessment, and improves reproducibility in ALS/FTD research.

## Materials and Methods

### Stem Cell Culture

The KOLF2.1J iPSC line has been previously described^22^. iPSCs were cultured in E8 Flex media (Gibco, cat# A2858501) on Vitronectin (Gibco, cat# A14700) coated plates and maintained at 37°C in 5% CO2 and hypoxic conditions. Cells were routinely passaged using Versene (Gibco, cat# 15040066) and Accutase (Sigma, cat# A6964) for CRISPR-Cas9 genome editing experiments.

### Induced NGN2 Lower Motor Neuron Differentiation

iPSCs were differentiated into lower motor neurons (liMNS) adapted from a previously published protocol (Figure 2A). Briefly, on day 0, iPSCs were plated at a density of 1 × 10⁶ cells per well in E8-Flex medium supplemented with 10 μM Y-27632 (Abcam, cat# ab120129) on Geltrex-coated 6-well plates. On day 1, cells were switched to induction medium composed of DMEM/F-12 (Gibco, cat# 10565018), supplemented with 1x GlutaMAX (Gibco, cat# 35050061), 1x N2 (Gibco, cat# 17502001), 1x NEAA (Gibco, cat# 11140050), 10 μM SB431542 (Cayman Chemicals, cat# 13031-1), 1 μM Dorsomorphin (Cayman Chemicals, cat# 11967), 1 μM EC23 (Cayman Chemicals, cat# 9002073), 1 μM SAG (Cayman Chemicals, cat# 912545-86-9), and 5 μM Y-27632 (Abcam, cat# ab120129). From days 2 to 3, induction medium was supplemented with 2 μg/ml doxycycline, and Y-27632 was omitted. From days 4 to 7, medium was replaced with Neurobasal Plus (Gibco, cat# A3582901), supplemented with 1x GlutaMAX, 1x N2, 1X B27 Plus (Gibco, cat# A3582801), 1XNEAA, 1 μM EC23, 1 μM SAG, 2 μg/ml doxycycline, and 10 ng/ml each of BDNF, GDNF, and CNTF (PeproTech, cat # 450-02, 450-10, 450-13). On day 7, cells were dissociated using Accutase and replated on poly-L-ornithine (Sigma, cat# P4957-50ML) and 20 μg ml−1 laminin (Sigma, cat# L2020) coated dishes in neuronal maturation medium composed of Neurobasal Plus, 1x GlutaMAX, 1x N2, 1x B27 Plus, 10 ng/ml BDNF/GDNF/CNTF, 5 μM Y-27632, and 1 μM Adarotene (MedChemExpress, cat# HY-14808). Cells were maintained in maturation medium with half media changes twice weekly, omitting Y-27632 and Adarotene after initial plating.

### Donor Plasmid and cloning

The *C9ORF72* HRE donor plasmid used was engineered via adaptation of a repeat construct originally designed for mouse genome targeting^42^, and contains 1027 G_4_C_2_ repeats flanked by 2 kilobases of human homology upstream and 3 kilobases downstream of the repeat locus, respectively, within the pJazz backbone^43^. Additionally, the plasmid contains a positive selection cassette, expressing mNeonGreen and conferring puromycin resistance, flanked by PiggyBac transposase cut sites (Supplementary Figure 1A) to allow for the seamless excision of the cassette following selection of iPSC colonies positive for the integration of the HRE.

The pJazz plasmid was transformed into BigEasy® TSA™ Electrocompetent Cells (Lucigen, cat# 60224-1) following the manufacturer’s instructions. Prior to transformation, the plasmid was denatured at 70 °C. Cells were thawed on ice and transferred to a chilled electroporation cuvette. 1 µL of denatured plasmid was added, and electroporation was performed using a Bio-Rad Gene Pulser under manufacturer-recommended settings. Following electroporation, recovery medium was added, and cells were incubated at room temperature with shaking for 2 hours. The transformation mixture was plated on LB Lennox-kanamycin agar and incubated at 25 °C for 72 hours. Individual colonies were picked and cultured overnight in Terrific Broth (Gibco, cat# A1374301). Plasmid DNA was isolated via miniprep, and clones were screened by restriction digest to confirm the integrity of the repeat region and absence of hexanucleotide repeat expansion (HRE) retraction. Genomic DNA was digested with EcoRI-HF (NEB, cat# R3101) and resolved on a 1% agarose gel. A 14.6 kb band indicated an intact pJazz plasmid containing the full-length G_4_C_2_ repeat sequence.

### gRNA design and cloning

The CRISPR approach followed a two-step strategy (Figure **1**A). First, a novel guide RNA (gRNA) recognition site was introduced upstream of the two G_4_C_2_ repeats in the KOLF2.1J iPSC line (Figure 1B). This insertion was designed to prevent cleavage of the pJazz donor plasmid containing 1027 G_4_C_2_ repeats.

The gRNA target sequence (ACTCAGGAGTCGCGCGCTAGGTTTTAGAGCTATGCT) was designed using IDT online software. The gRNA with the highest predicted on-target efficiency and lowest off-target potential was selected to generate the *C9ORF72* intermediate line. This gRNA targets a region upstream of the two endogenous G_4_C_2_ repeats in the *C9ORF72* locus of the KOLF2.1J line (Supplementary Table 1). To introduce the novel cut site, a ssODN (IDT) was designed with 40 bp of homology flanking the intended mutation site (Supplementary Table 1). The endogenous *C9ORF72* sequence GTCGC**<u>GCGC</u>**TAG**<u>GG</u>**GCCGGGGCC to GGAGTCGC**<u>ATACG</u>**TAG**<u>TG</u>**CCGGGGCC was first mutated using a Cas9:deadCas9 approach to generate the heterozygous intermediate line. iPSC colonies were screened for the insertion of the new mutations by Sanger sequencing (Figure 1B). TOPO TA cloning was undertaken to ensure heterozygosity and additional quality control checks to ensure the absence of off-target insertion or deletions (Supplementary Figure 3B).

Following successful integration of the ssODN sequence into the *C9ORF72* locus, a second gRNA (GAGTCGCATACGTAGTGCCGGTTTTAGAGCTATGCT) was designed to target the novel cut site while preserving the integrity of the pJazz plasmid and homology arms. Subsequently, the same gRNA was used to revert the novel gRNA target site to the wild-type *C9ORF72* sequence, thereby generating a CRISPR control iPSC line. This ensured the control line underwent the same number of genome editing steps as the *C9ORF72* knock-in lines, minimizing the likelihood of clonal selection bias and unintended indel generation.

### Genome Editing via CRISPR/Cas9

Healthy iPSCs were cultured to ∼80% confluency prior to CRISPR-mediated genome editing. The gRNA was prepared by annealing crRNA and tracrRNA (IDT) at 95 °C for 5 minutes. The ribonucleoprotein (RNP) complex was formed by incubating the gRNA with Cas9 (IDT, Alt-R® S.p. HiFi Cas9 Nuclease V3, cat# 1081060) and dCas9 (IDT, Alt-R® S.p. dCas9 Protein V3, cat# 1081066) at a molar ratio of 1:1.5 for 20 minutes at room temperature. iPSCs were dissociated using Accutase, pelleted at 300*×g* for 4 minutes, and washed once with PBS. Cell pellets were resuspended in 20 µL of Nucleofector Solution (Lonza, P3 Primary Cell 4D-Nucleofector, cat# V4XP-3032). The RNP complex was added to the cell suspension along with 2 µg of ssODN and electroporation enhancer (IDT, Alt-R™ Cas9 Electroporation Enhancer, cat# 1075915). Cells were nucleofected using a Lonza 4D-Nucleofector (cat# AAF-1003B), then plated in E8 Flex medium supplemented with 10 µM Y-27632 and HDR enhancer (IDT, cat# 10007910). After recovery, cells were dissociated to single cells and sparsely replated in a D150 dish. Single iPSC cells were allowed to grow until they reached a colony size of >∼50 cells per colony. These colonies were manually picked and expanded. Genomic DNA was isolated using the Qiagen DNeasy Blood and Tissue Kit (Qiagen, cat# 69504), and clones were screened for correct insertion of the 5 bp mutation using Sanger sequencing and to assess both on- and off-target modifications.

### Insertion of G_4_C_2_ Hexanucleotide Repeat Expansion

A pJazz plasmid harbouring 1027 G_4_C_2_ repeats was used to target the endogenous *C9ORF72* locus in a heterozygous intermediate line. The gRNA was assembled as above and incubated with Cas9 (IDT, cat# 1081060) for 20 minutes to form the RNP complex. iPSCs were dissociated with Accutase, pelleted, and resuspended in 20 µL of Nucleofector Solution. The RNP complex, 600 ng of the pJazz plasmid, and electroporation enhancer (IDT, cat# 1075915) were added to the cells, which were nucleofected and plated onto Vitronectin-coated 6-well plates in E8 Flex medium supplemented with 10 µM Y-27632 and HDR enhancer. After 48 hours, cells were selected with 1 µg/mL puromycin. Colonies showing nuclear mNeonGreen expression were screened by PCR using primers flanking the integration site (Supplementary Table 2). Positive clones were subcloned via single-cell sorting to ensure clonality, and repeat expansion was verified using repeat-primed PCR. Fluorescence-activated cell sorting (FACS) was used to isolate mNeonGreen-positive populations.

### Removal of the pJazz Selection Cassette

After successful HDR-mediated insertion of the repeat expansion, the selection cassette was excised using a hyperactive, excision-only PiggyBac transposase (iPB7^R372A/K375A/D450N^) ^25^.The expression plasmid for the hyperactive excision-only PiggyBac transposase was created by cloning a codon optimised cDNA (Geneart, Life Technologies) into the EcoRI and XbaI sites of pRR-EF1a-Puro under the EF1alpha promoter^24^. Clones confirmed to harbour the repeat were nucleofected with this plasmid. Cassette removal was validated via PCR with primers spanning the selection cassette junctions (Supplementary Table 2).

### Insertion of the Doxycycline Inducible-NGN2 Cassette

A doxycycline-inducible *NGN2* expression cassette (excision-TO-hNGN2, Addgene, cat# 172115) was inserted via PiggyBac transposition. iPSCs were cultured to ∼80% confluency and dissociated using Accutase. Approximately 2×10⁶ cells were resuspended in 20 µL Nucleofector Solution and nucleofected with 3 µg of total DNA (NGN2 cassette and PiggyBac transposase plasmid, 3:1 ratio). Following recovery, FACS was used to isolate BFP-positive cells.

### TOPO TA Cloning for Heterozygous Clone Selection

TOPO TA cloning was used to identify heterozygous intermediate clones prior to pJazz insertion. PCR was performed on 100 ng genomic DNA using standard Taq polymerase (1× buffer, 50 mM dNTPs, 150 ng primers). PCR products with 3′ A-overhangs were cloned using the TOPO TA cloning kit (PCR product 4 µL, salt solution 1 µL, TOPO vector 1 µL) and incubated for 5 minutes at room temperature. 2 μl of the reaction were transformed into One Shot® chemically competent E. coli cells. After a 30-minute incubation on ice and a 30-second heat shock at 42 °C, cells were returned to ice, and 600ul of S.O.C. medium was added, and samples shaken at 37 °C for 1 hour. 50 μl of the transformation were plated onto LB-carbenicillin plates and incubated overnight. Colonies were screened for heterozygous insertion via colony PCR and sequencing.

### Off-Target Analysis

All edited iPSC clones underwent off-target analysis to confirm genomic integrity. The top three potential off-target loci for each gRNA were identified using IDT’s gRNA Design Checker tool. Genomic DNA was extracted, and each locus was PCR-amplified and compared against the parental KOLF2.1J line via Sanger sequencing to confirm the absence of indels or insertions. Target sites and primers are listed in Supplementary Table 3. Karyotyping of all cell lines was undertaken using the Karyostat+ chromosomal microarray service from Thermo Fisher. All cell lines were karyotypically normal Supplementary Figure 3A.

### Repeat prime PCR

Repeat prime PCR was performed as previously described^44^. Briefly, 100 ng of genomic DNA was amplified using the 1× Roche FastStart Master Mix (Roche, cat# 4710436001) supplemented with 0.9 mM Betaine, 7% DMSO, 0.9 mM MgCl₂, and 0.18 mM 7-deaza-dGTP. PCR products were mixed with HiDi Formamide (ABI, cat# 4311320) and LIZ500 size standard (ABI, cat# 4322682) and analysed using the ABI 3730 DNA Analyzer. Data were visualised using Thermo Fisher Peak Scanner software (apps.thermofisher.com).

### Southern Blotting

Genomic DNA was extracted from the isogenic *C9ORF72* iPSC lines using the Qiagen DNeasy Blood & Tissue Kit (Qiagen, cat# 69504) as per the manufacturer’s protocol. Southern blot analysis of the repeat size was undertaken as previously described^1^ with some minor modifications. Genomic DNA was digested with Alu1/DdeI overnight at 37°C and 12 µg was electrophoresed on a 0.8% agarose gel alongside the DIG-labelled DNA molecular weight marker VII (Sigma-Aldrich, cat# 11669940910). The DNA was transferred overnight to a positively charged nylon membrane (Sigma-Aldrich, cat #1141724001) by capillary blotting and was crossed-linked the following morning by UV. The membrane was prehybridized in Roche DIG Easy Hyb solution (Sigma-Aldrich, cat# 11796895001) with the addition of 100 µg/ml denatured salmon sperm (Thermo, cat# 15632011) for 3 hours at 48°C. The membrane was probed overnight at 48°C with 10ng/ml of a DIG labelled oligonucleotide comprising of five hexanucleotide repeats (GGGGCC)_5_ (IDT). Following hybridization, the membrane was washed in a 2x sodium citrate (SSC) and 0.5% sodium dodecyl sulphate for 15 minutes while the hybridization oven ramped from 48°C to 65°C. The membrane was then washed in 2x SSC and 0.5% SDS for 15 mins, followed by 15 min washes in 0.5X SSC and 0.5% SDS and 0.15X SSC and 0.5% SDS. Antibody detection was carried out following the DIG Application Manual protocol (Roche Applied Science) using the DIG wash and block buffer set (Roche, cat# 11585762001) and ready-to-use CSPD (Sigma-Aldrich, cat# 11755633001). Signals were visualised using ChemiDoc XRS+ system (BioRad). Membranes were exposed for 45 mins with images collected every 4 mins. Hexanucleotide repeat expansions were sized using Image Lab 6.1 software (BioRad) by comparison to DIG-labelled DNA molecular weight marker VII and subtraction of the genomic non repeat region of *C9ORF72* flanking the AluI/DdeI restriction cut sites (156bp).

### Fluorescence In-Situ Hybridisation

RNA-FISH was used to detect *C9ORF72* sense and antisense RNA foci following previously established protocols^31^. At DIV7, 25,000 liMNs were seeded into 96-well PhenoPlates (Revvity, cat# 6055300) pre-coated with poly-L-ornithine and laminin and maintained until DIV 28 (N = 3 differentiations per line, 2 technical replicates each). Cells were fixed in 4% paraformaldehyde (PFA) for 20 min, then dehydrated through 70% and 100% ethanol and stored in 100% ethanol at -80°C. Before hybridisation, cells were rehydrated and incubated in pre-hybridisation buffer (40% formamide, 10% dextran sulfate, 2 mM Ribonucleoside Vanadyl Complex, 2× SSC) for 5 min at room temperature (RT), then permeabilised in 0.2% Triton X-100 in PBS for 10 min. Hybridisation was performed overnight at 66°C (sense) or 60°C (antisense) with 40 nM TYE563-labelled LNA probes (Qiagen): sense (5′-CCCCGGCCCCGGCCCC), antisense (5′-GGGGCCGGGGCCGGGGCC). After hybridisation, cells were washed and immunolabelled with rabbit anti-βIII Tubulin (1:500, A85363) overnight at 4°C, followed by goat anti-rabbit Alexa Fluor 488 secondary antibody. Nuclei were stained with DAPI (100 ng/mL) prior to image acquisition.

### Whole Genome Sequencing

Cell pellets were collected from all iPSC lines. Genomic DNA was extracted, and sample QC was undertaken at Novogene UK. Whole genome sequencing of all samples was performed on an NovaSeq X Plus Series (PE150), producing 354.8 – 367.8 million pairs of 150bp reads per sample. Reads were aligned using bwa mem v0.7.17-r1188 to a custom reference consisting of the full pJAZZ plasmid sequence appended to the GRCh38 human genome. Several default bwa parameters were relaxed (-M -Y -k 13 -L 4,4 -U 10 -T 12 -a) to aid in the mapping and identification of chimeric reads and read pairs. Sorted and indexed BAM files were viewed in the Integrative Genomics Viewer v2.4.7.

### Nanopore Adaptive Long-Read Sequencing

Genomic DNA was extracted using the Monarch HMW DNA Extraction Kit for Cells and Blood (New England BioLabs, T3050) following the manufacturer’s instructions. The DNA quality, quantity and size distribution were assessed using a Qubit fluorometer with the Qubit dsDNA BR Assay Kit (Thermo Fisher Scientific, Q32850) and a Femto Pulse system with the gDNA 165 kb analysis kit (Agilent Technologies, FP-1002-0275).

A protocol for sample preparation and Nanopore processing was developed by the Genomics England Scientific R&D Team with some minor modifications. 6-6.4 µg of DNA was fragmented to 10-15 kb using Covaris g-TUBEs (520079, Covaris). Samples were centrifuged using an Eppendorf 5424 (022620401, Eppendorf) centrifuge for 1 minute at 4500-4800 rpm, rotated 180 degrees and centrifuged for a further minute at 4500-4800 rpm. Fragments under 5 kb were reduced using the SRE XS kit (SKU 102-208-20,0 PacBio). An equal volume of Buffer SRE-XS was added to the fragmented DNA sample and centrifuged at 10,000 *xg* for 30 minutes before discarding the supernatant. The DNA pellet was washed with 200 µL of 70% ethanol for a total of two ethanol washes. The remaining ethanol was evaporated off at 37°C for 2-15 minutes as required. 50 µl of PacBio Buffer EB was used to resuspend the pellet at 37°C for 10-20 minutes then 4°C overnight. For library preparation 1-1.5 µg of sample in 48 µL of nuclease-free water (NFW) was mixed with 3.5 µL NEBNext FFPE DNA Repair Buffer, 3.5 µL Ultra II End-Prep Reaction Buffer, 2 µL NEBNext FFPE DNA Repair Mix (M6630, NEB) and 3 µL Ultra II End-Prep Enzyme Mix (E7546, NEB) and incubated at 20°C for 10 minutes followed by 65°C for 10 minutes. The reaction was then incubated at room temperature for 10 minutes on a hula mixer with 60 µL of AMPure XP Beads (A63881, Beckman Coulter). The beads were pelleted on a magnet and washed with 200 µl of 70% ethanol twice before eluting in 64 µL NFW at room temperature for 10-30 minutes. The library was processed using Oxford Nanopore Technologies (ONT) Ligation Sequencing Kit V14 (SQK-LSK114). 62 µL of eluted DNA was added to a mix of 25 µL ONT Ligation Buffer, 8 µL NEBNext Quick T4 DNA Ligase (E6056, NEB) and 5 µL ONT Ligation Adapter and incubated at room temperature for 30 minutes. 40 µL of AMPure XP beads were added and incubated on a hula mixer at room temperature for 10 minutes then pelleted on a magnet. The beads were washed twice with 250 µL ONT Long Fragment Buffer then eluted in 34 µL ONT Elution Buffer at 37°C for 15 minutes. 10-15 fmol of library was loaded onto a single PromethION Flow Cell (R10.4.1) following manufacturer’s instructions. Adaptive sampling was carried out as described below. The run lasted 72 hours with two 1-hour nuclease flushes between 20-24 hours and 44-48 hours using the ONT Flow Cell Wash Kit (EXP-WSH004) following manufacturer’s instructions. The library was stored at 4°C during the run and 10-15 fmol of library was loaded after each nuclease flush. For adaptive sampling and analysis, all code can be found at: https://github.com/rainwala/GIU-rep_exp_blast_finder. Adaptive Sampling was carried out using the Adaptive Sampling option on MinKnow version 24.02.10. The reference genome used was GRCh38.p14 (https://www.ncbi.nlm.nih.gov/datasets/genome/GCF_000001405.40/) and the co-ordinates were specified in bed format as follows: chr9 27543537 27603536 C9ORF72_expansion_region. This corresponds to the file C9ORF72_adaptive.bed in the Github repository above. Once sequencing was complete, the nextflow process rep_exp_blast_finder.nf was run on the sequencing output to generate the final output.

### Removal of Cassette Integration Verification

Successful removal of the selection cassette at the endogenous *C9ORF72* locus was confirmed by PCR, whole-genome sequencing (WGS), and Nanopore adaptive long-read sequencing. WGS detected a partial plasmid backbone insertion downstream of the HRE insertion within intron 1a- to our knowledge a region not implicated in RAN translation or transcriptional activity. No chimeric or off-target integrations were observed, confirming insertion at the correct locus. Nanopore sequencing, which provides higher accuracy across repetitive regions, verified the absence of the pJazz selection cassette and confirmed seamless integration of the HRE and cassette removal. The minor plasmid sequence detected by WGS likely reflects residual contamination or a minor mixed population. Overall, long-read data confirm the integrity and correct configuration of the *C9ORF72* locus following cassette excision.

### RT-qPCR

Total RNA was extracted from iPSCs and DIV 28 liMNs cell pellets using the Reliaprep RNA miniprep system (Promega, cat# Z6010) as per the manufacturers protocol. A total of 500-1mg of RNA was used to synthesize cDNA from each sample using the LunaScript® RT SuperMix following the manufacturers protocol. Quantitative qPCR was perform using PowerUp™ SYBR™ Green Master Mix (Applied Biosystems, cat# A25742) for all targets aside from *C9ORF72* variant 2 which was measured using the TaqMan™ Fast Advanced Master Mix (Applied Biosystems, cat# 4444556). Samples were run in triplicate with 7.5ng of cDNA and primers at a final concentration of 600nM per reaction and signal was measured using the QuantStudio7 (Thermo Scientific). Quantitative levels for all genes assayed was normalised to the signal of *GAPDH*. All experiments were performed using the two control lines (KOLF2.1J, WT Rev) compared to the two clonal lines containing the HRE (C06, C14) with 3 independent biological replicates (differentiations) and 3 technical replicates. Transcripts were amplified using the following primers; *GAPDH* (agtcagccgcatcttctttt, accagagttaaaagcagccc)^32^, *MNX1* (catgatcctgcctaagatgcc, cgacaggtacttgttgagctt)^32^, *OCT3/4* (atgcattcaaactgaggtgcctgc, aacttcaccttccctcgaacgagt), *C9ORF72* total (actggaatggggatcgcagca, acctgatcttccattctctctgtgcc)^30^, *C9ORF72* Variant 2 (gcggtggcgagtggatat, tgggcaaagagtcgacatca, /56-FAM/ATTTGGATA/ZEN/ATGTGACAGTTGG)^28^, *C9ORF72* Intron 1a (ccccactacttgctctcaca, ggttgtttccctccttgtt) ^25^.

### Western Blotting

Protein was extracted from cell pellets in RIPA buffer (2% SDS + protease inhibitors) and sonicated (Bioruptor, Diagenode) at 4°C (3 × 10 s). Lysates were centrifuged at 16,000 rpm for 20 min at 16°C. Protein concentration was determined using the Pierce BCA assay (Thermo Scientific, cat# 23225). Equal protein volumes were loaded onto NuPAGE Bis-Tris Mini Gels (Invitrogen, cat# NP0321BOX), transferred to nitrocellulose membranes (cat# IB23002) using iBlot 2™. Membranes were blocked in 5% milk/TBS-T (0.1% Tween-20) and probed overnight at 4°C with anti-C9ORF72 (Genetex, GT1553-RB, 1:1000) and anti-GAPDH (Santa Cruz, sc32233, 1:1000) antibodies. IRDye® secondary antibodies were applied for 1h at room temperature. Blots were visualised using the Odyssey CLx Imager (LicorBio) and analysed using Image Studio.

### Immunocytochemistry

iPSCs and liMNs were cultured in 96-well PhenoPlates (Revvity, cat# 6055300). Cells were fixed in 4% PFA for 15 min at room temperature, washed with PBS, and blocked/permeabilised in 6% BSA + 0.5% Triton X-100 (TBS) for 30 min. Cells were incubated with primary antibodies (6% BSA + 0.1% Triton X-100 in TBS) overnight at 4°C: mouse anti-SSEA-4 (sc21704, 1:500), mouse anti-TRA-1-81 (sc21706, 1:100), rabbit anti-Nanog (sc33759, 1:200), Alexa Fluor® 568 Anti-Islet 1 (EP4182, 1:500), chicken anti-MAP2 (A85363, 1:500). Secondary antibodies (goat anti-mouse 488, anti-rabbit 555, anti-chicken 633, all 1:1000) were applied for 1h at room temperature. DAPI (100 ng/mL) was used to stain nuclei.

### MSD Immunoassay

Poly(GA) and poly(GP) levels were assessed via Meso Scale Discovery (MSD) immunoassay as previously described^45,46^. All DPR protein extracts were collected in RIPA buffer (2% SDS + protease inhibitors) and normalised to 0.5 mg/ml. 25 µl of each samples loaded in duplicate. Primary antibodies included anti-poly(GA) (MABN889, Sigma) and anti-poly(GP) (GP658, custom-made from Eurogentec), and detection used biotinylated antibodies (GA 5F2*, a kind gift from Dieter Edbauer and GP658*, custom-made from Eurogentec). Plates were read using MSD Sector Imager 2400.

### Image Acquisition and Analysis

Images were captured on a Nikon Eclipse Ti Inverted Spinning Disk Confocal microscope using 20× and 60× oil immersion objectives. For RNA-FISH, Z-stacks (0.15 µm interval) were acquired. Image quantification, including RNA foci and ISLET-1 staining, was conducted using NIS Elements 3D analysis pipelines using the same settings for all samples.

### Statistics & Reproducibility

Independent differentiations were considered biological replicates (represented as individual data points on graphs); replicate wells were technical replicates. A minimum of three biological replicates was included per experiment unless otherwise specified. Wells with detached or degenerated neurons were excluded from analyses. Statistic difference was tested by one-way ANOVA with post hoc Šídák’s multiple comparisons test with comparison versus controls using GraphPad Prism.

## Declarations

### Ethics approval and consent to participate

The research involves human iPSC lines, derived from the KOLF2.1j background obtained from Jackson laboratories, which has well-documented origin obtained under informed donor consent and under relevant ethical oversight^21,22^.

### Consent for publication

N/A

### Data Availability

All data generated and analysed during this study are included in this article and its supplementary information files, or available from the corresponding authors on reasonable request.

### Authors’ contributions

Conceptualisation of study, supervision, project administration, funding: SM, MDR. Methodology and study design: SM, MDR, NOB, EH. Generation of the linear pJazz plasmid: TC, RRN. CRISPR/Cas9 experiments and generation of the final NGN2-HRE lines and quality control: NOB, EH, STa. Characterisation of the cell line phenotypes: NOB, EH, DV. Repeat prime PCR: AJC, AMI. DPR MSD assay: MC, AMI. FISH analysis of RNA foci: NOB. Whole Genome Sequencing: STo. Nanopore sequencing: NOB, AA, KS, CFW. Formal analysis: NOB, SM, STo. Writing of original draft: NOB, SM, MDR, preparation of figures: NOB. Reviewing of draft: NOB, EH, RRN, AJC, MC, DV, STa, KS, ARA, STo, AI, AMI, TJC, SM, MDR.

## Acknowledgements

We thank Dr George Chennell and the team at the Wohl Cellular Imaging Centre at King’s College London for help with light microscopy and training. We thank Prof William C. Skarnes for sharing the KOLF2.1J cell line with us ahead of publication. BioRender was used for the creation of images in Figures 1-3.

## List of abbreviations

ALS: amyotrophic lateral sclerosis
ASO: antisense oligonucleotide
DPR: dipeptide repeat
FTD: frontotemporal dementia
HRE: hexanucleotide repeat
iPSC: induced pluripotent stem cell
liMNs: lower induced motor neurons
RAN: repeat associated non-ATG translation
rpPCR: repeat primed PCR
ssODN: single-stranded oligodeoxyribonucleotide

