## Supplementary Data for "CRISPR/Cas9-mediated generation of a *de novo C9ORF72* knock-in isogenic iPSC cell bank to model ALS and FTD"

### Supplementary Figures

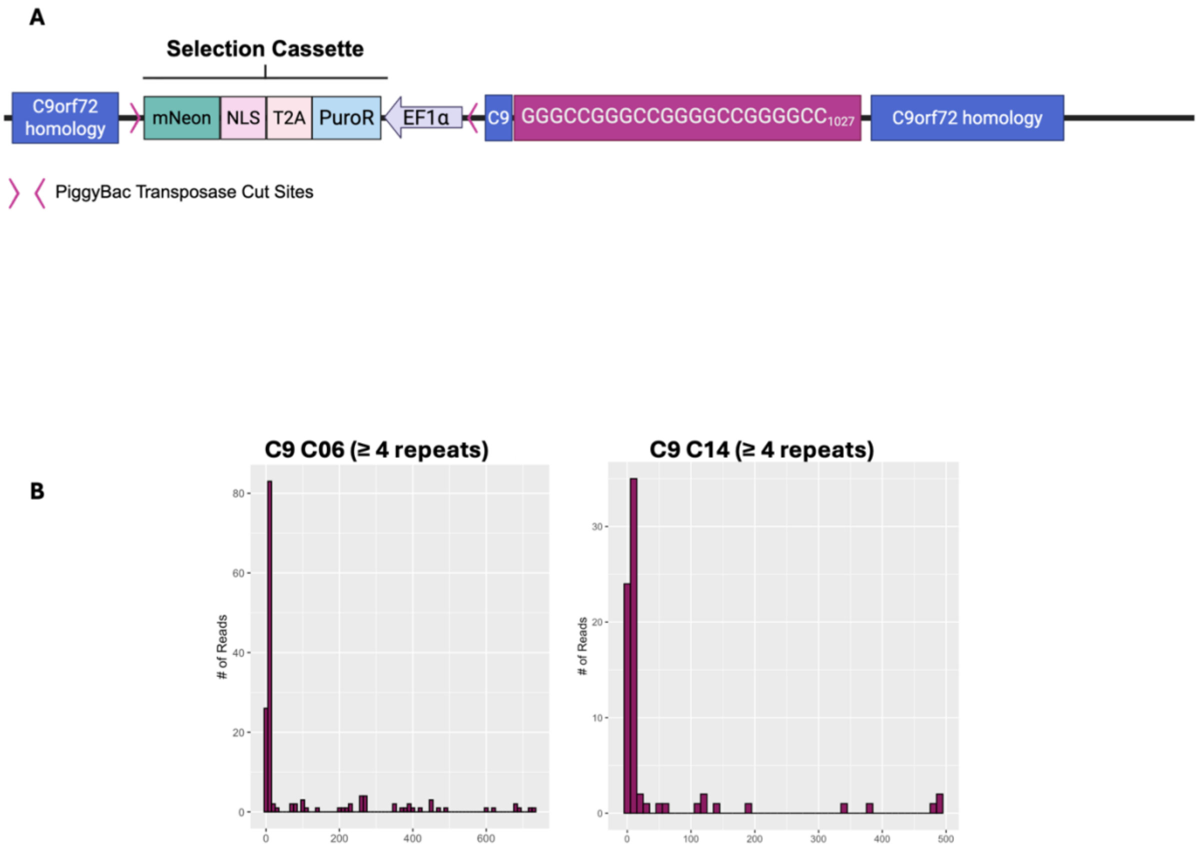


**Supplementary Figure 1:** (A) The linearised pJazz plasmid was constructed to contain 2kb of homology to the *C9ORF72* gene, a positive selection cassette, 1027 $G_4C_2$ repeats and 2.8kb of *C9ORF72* homology downstream of the HRE. The positive selection cassette consisted of an mNeonGreen-NLS (nuclear localisation sequence) reporter and a puromycin resistance cassette and was flanked by 2 PiggyBac transposase cut sites. (B) Distribution of reads from Nanopore sequencing data which contained over 4 $G_4C_2$ aligned to the endogenous *C9ORF72* locus.

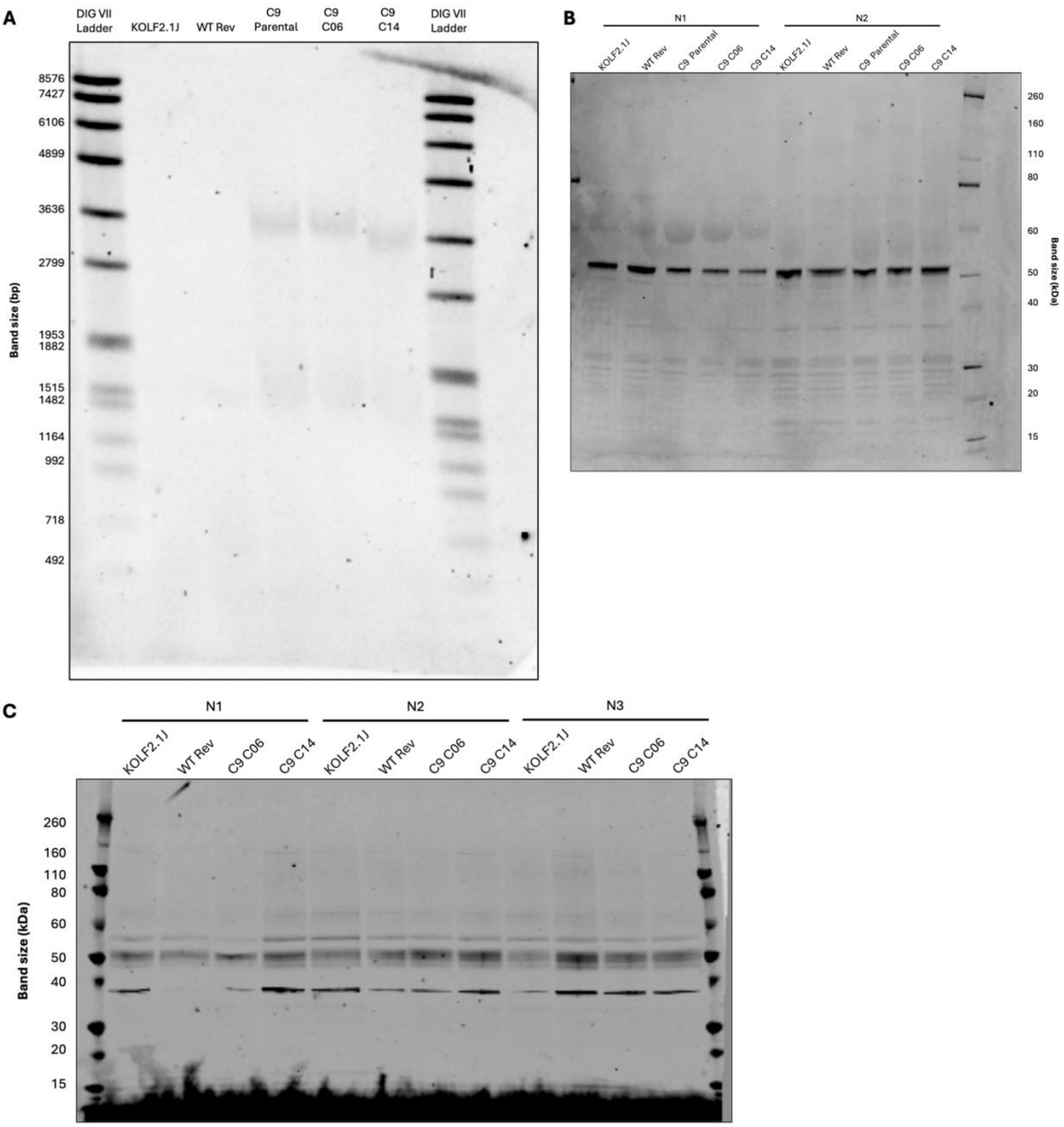


**Supplementary Figure 2:** (A) Full southern blot showing samples run against DIG VII ladder. (B) Full western blot of C9ORF72 protein levels in iPSC lines. (C) Full western blot of C9ORF72 protein levels in DIV 28 liMNs.

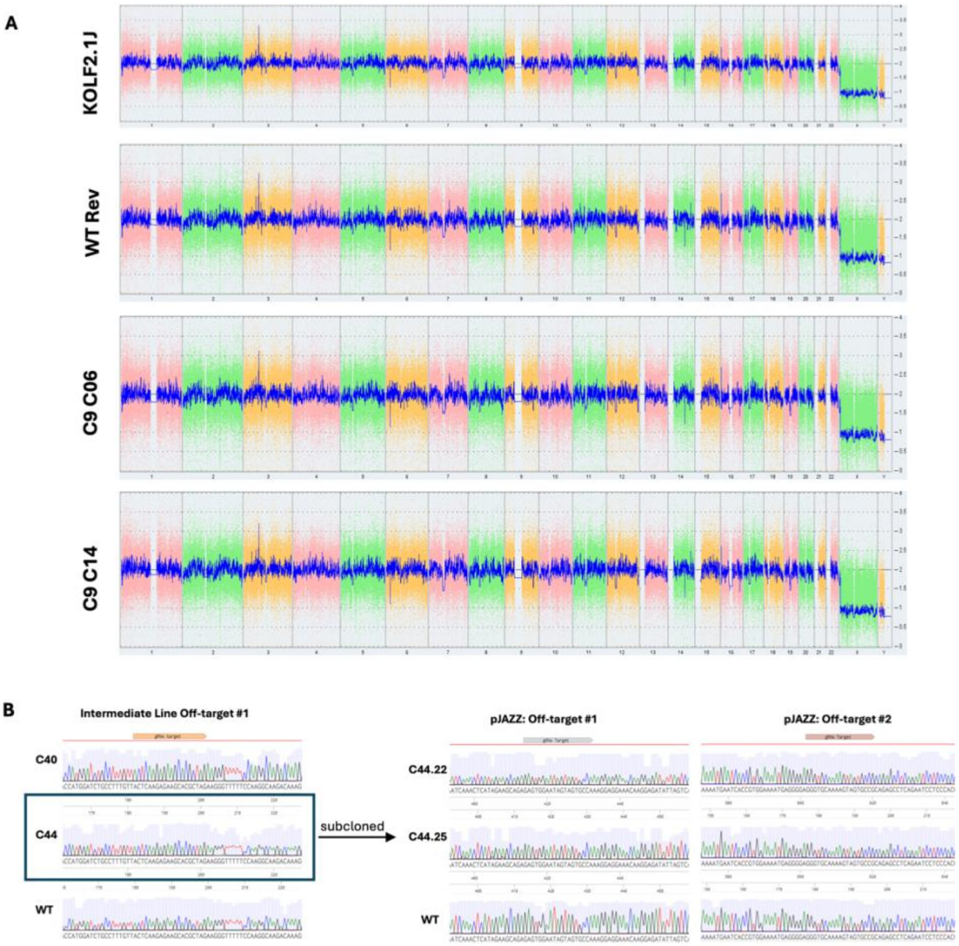


**Supplementary Figure 3:** Quality control for genomic integrity in the HRE derived lines and wildtype controls. (A) Karyostat analysis demonstrates a normal cellular karyotype in all cell lines. (B) Off-target analysis of genomic regions following CRISPR/Cas9. C44 is the intermediate landing pad line which has no in/dels present in the top 3 off-target regions. A second guide was then used for the generation of the WT Rev and HRE knock-in lines. Neither of these lines have off-target indels in the top three predicted targets.

### Supplementary Tables

#### Supplementary Table 1

| Oligonucleotide | Sequence-IDT |
|---|---|
| Intermediate ssODN | /AlT-R-HDR1/C*C*GCCCCGGGCCCGCCCCCGGGCCCGCCCCGACCACGCCCCGGCCCCGGCACTACGTATGCGACTCCTGAGTTCCAGAGCTTGCTACAGGCTGCGGTTGTTTCCCTC*C*T/AlT-R-HDR2/ |
| Intermediate gRNA | /AlTR1/rArCrUrCrArGrGrArGrUrCrGrCrGrCrGrCrUrArGrGrUrUrUrUrArGrArGrCrUrArUrGrCrU/AlTR2/ |
| WT Rev ssODN | /Alt-R-HDR1/C*C*CGCCCCCGGGCCCGCCCCGACCACGCCCCGGCCCCGGCCCCTAGCGCGCGACTCCTGAGTTCCAGAGCTTGCTACAGGCTGCGGTT*G*T/Alt-R-HDR2/ |
| gRNA pJazz/ WT Rev gRNA | /AlTR1/rGrArGrUrCrGrCrArUrArCrGrUrArGrUrGrCrCrGrGrUrUrUrUrArGrArGrCrUrArUrGrCrU/AlTR2/ |

Guide RNA oligonucleotide and ssODN sequences used for CRISPR/Cas9

#### Supplementary Table 2

| Region | | Primers | Sequence |
|---|---|---|---|
| PCR 1 | Genome- pJazz Cassette | F | gcagctggacttggagagaattacactgtggg |
| | | R | cattgacaagcacgcctcac |
| PCR 2 | PB 5'LTR | F | aagagcaggtgtgggtttag |
| | | R | gcatggacgagctgtacaag |
| PCR 3 | mNeonGreen- Puromycin Resistance | F | gacacctactcagacaatgc |
| | | R | agtgggtggagactgaagttag |
| PCR 4 | Puromycin resistance- EF1α promoter | F | ggaggccttccatctgttg |
| | | R | gtgcagtagtcgccgtgaac |
| PCR 5 | EF1α promoter- upstream of HRE insert | F | gactacccccgtccgattctcg |
| | | R | ggctgcggttgtttccctcctt |

Primers used to determine whether integration and removal of the positive selection was successful during CRISPR/Cas9

**Supplementary Table 3**

| Line | gRNA Seq | On-target score | Off-target score | Chromosome target | Forward Primer | Reverse Primer |
|---|---|---|---|---|---|---|
| Intermediate | actcaggagtc gcgcgctag | 15 | 96 | chr12:+72736949 | aaaattctctggcccctgc | gctgagtgttcttgttcccc |
| | | | | chr17:-8470914 | tgcacggctaattcatggtg | gagccaagatcacgccattg |
| | | | | chr11:-2700847 | gttctctgcgtgatgtgttca | ggcagtgaagatgaatggc c |
| *C9ORF72* HRE/WT Rev | gagtcgcatac gtagtgccg | 45 | 92 | chr3:+105216279 | agtgtacaagtctttcacctcct | tggaaagcatcaccacaatc a |
| | | | | chr20:+57069832 | ggacaggaacaaggacagg a | ccgtgttgttcagtgtggaa |
| | | | | chr14:-87081200 | tcaaggtgacaggtgcttca | agctaagtttcttcttcaggact |

Primer sequences used to ensure the absence of off-target editing

**Supplementary Table 4**

| Target | Primer Sequence |
|---|---|
| *C9ORF72* intron 1a-repeat | CCCCACTACTTGCTCTCACA |
| | CGGTTGTTTCCCTCCTTGTT |
| *C9ORF72* Total | ACTGGAATGGGGATCGCAGCA |
| | ACCTGATCTTCCATTCTCTCTGTGCC |
| *C9ORF72* variant 2 | GCGGTGGCGAGTGGATAT |
| | TGGGCAAAGAGTCGACATCA |
| | /56-FAM/ATTTGGATA/ZEN/ATGTGACAGTTGG |

qPCR primers sequences used for analyzing *C9ORF72* isoform expression in *C9ORF72* HRE knock-in and control iPSCs and liMNs
